# Deletion of *cftr* leads to an excessive neutrophilic response and defective tissue repair in a zebrafish model of sterile inflammation

**DOI:** 10.1101/842195

**Authors:** Audrey Bernut, Catherine A. Loynes, R. Andres Floto, Stephen A. Renshaw

## Abstract

Inflammation-related progressive lung destruction is the leading causes of premature death in cystic fibrosis (CF), a genetic disorder caused by mutations in the cystic fibrosis transmembrane conductance regulator (CFTR) gene. However, therapeutic targeting of inflammation has been hampered by a lack of understanding of the links between a dysfunctional CFTR and the deleterious innate immune response in CF. Herein, we used CFTR-depleted zebrafish larvae as an innovative *in vivo* vertebrate model, mimicking aspects of the inflammatory pathology of CF-related lung, to understand how CFTR dysfunction leads to abnormal inflammatory status in CF.

We show that impaired CFTR-mediated inflammation correlates with an exuberant neutrophilic response after injury: CF zebrafish exhibit enhanced and sustained accumulation of neutrophils at wounds. Excessive epithelial oxidative responses drive enhanced neutrophil recruitment towards wounds. Persistence of neutrophils at inflamed sites is associated with impaired reverse migration of neutrophils and reduction in neutrophil apoptosis. As a consequence, the increased number of neutrophils at wound sites causes tissue damage and abnormal tissue repair. Importantly, the pro-resolution molecule Tanshinone IIA successfully re-balances inflammation both by accelerating inflammation resolution and by improving tissue repair in CFTR-deficient animal.

Larval zebrafish giving a unique insight into innate immune cell function in CFTR deficiency, our findings bring important new understanding of the mechanisms underlying the inflammatory pathology in CF, which could be addressed therapeutically to prevent inflammatory lung damage in CF patients with potential improvements in disease outcomes.

## Introduction

Cystic fibrosis (CF) is a genetic disease resulting from mutations in the cystic fibrosis transmembrane conductance regulator (CFTR) and causes premature death by progressive respiratory failure, itself caused by lung destruction from a vicious circle of repetitive infection and excessive inflammation *(1, 2)*. It is commonly assumed that the CF-related lung pathology is primarily an infectious disorder: susceptibility to invading pathogens results from airway mucus obstruction and collapse of mucociliary clearance and that the resultant persistent infection drives chronic inflammatory lung damage *(3)*. However, in CF airways, there is an abnormal inflammatory phenotype often present in the absence of detectable infection *(4)*, suggesting that CFTR dysfunction might also cause primary defects in lung innate immunity, leading to an early pro-inflammatory state and raising the question of what drives non-infectious inflammation in CF.

So far, previous attempts to define a direct role for CFTR in host innate immune potential, using patient derived cells *(5)* or mammalian models *(6)*, have yielded contradictory results and have not been able to dissociate direct effects of CFTR dysfunction from the consequences of chronic inflammation on cellular function. Consequently, the mechanisms by which CFTR directly regulates innate immunity and how CF mutations contribute to inflammatory pathogenesis in CF have remained obscure. There is therefore a pressing need to develop tractable *in vivo* models allowing direct observation of CFTR-dependent effects on innate immune responses in the absence of a pre-existing inflammatory environment.

Zebrafish innate immunity is closely homologous to that of humans *(7)*, while their optical transparency allows non-invasive, real-time monitoring of inflammatory processes in the whole organism *(8)*. Therefore, zebrafish innate immune cell behaviour and function can be readily visualised at sub-cellular resolution, allowing new understanding of innate immune potential in inflammation. In particular, zebrafish have emerged as a powerful model to investigate inflammatory processes, with the ability to recapitulate many aspects of human inflammatory disease *(9)*. Remarkably, zebrafish CFTR retains close sequence identity with human (56.24% identity) *(10, 11)*. Moreover, like mammals, zebrafish CFTR is expressed in epithelial surfaces and myeloid cells *(12, 13)* and plays an important role in homeostatic balance of fluid composition *(14, 15)*. Several phenotypes that mirror many aspects of human CF were also reported in CFTR-defective zebrafish, including pancreatitis *(16)*, anaemia *(17)* and increased susceptibility to CF bacterial pathogens, such as *Pseudomonas aeruginosa (12)* and recently *Mycobacterium abscessus* complex *(13)*. The high level of genetic and functional conservation between the zebrafish and mammalian CFTR and innate immune systems, as well as the lack of a pre-existent CF-related inflammatory phenotype, make zebrafish a clinically-relevant system to investigate immune pathophysiology in CF and could provide insights not available in existing models.

While the absence of lungs might at first sight appear to reduce the translational relevance of this model, our data suggest that changes to CFTR function in epithelial and myeloid cells are conserved across tissues and species. Exploiting CFTR-depleted zebrafish larvae as an innovative vertebrate organism, we show here that *i)* there is a direct role for CFTR function in regulation of host innate immunity and tissue repair, *ii)* the inflammatory pathology of CF is determined by alterations in neutrophil apoptosis and reverse migration, and finally, *iii)* the pro-resolution molecule, Tanshinone IIA, can restore normal levels of inflammation and tissue repair in CF, with important implications for the treatment of CF.

## Results

### Loss of CFTR function promotes overactive neutrophilic inflammation to sterile tissue injury

To assess the role of CFTR in regulating host inflammatory potential *in vivo*, expression of *cftr* was knocked-out using CRISPR-Cas9 technology and/or knocked-down by morpholino-modified antisense oligonucleotides *(13)* (Fig. S1, A and B). In zebrafish larvae, tail amputation triggers leukocyte infiltration towards the wound, accurately mimicking the kinetics seen in mammalian inflammatory responses *(18–20)*. At these early stages, there is no infectious pathology in routine husbandry conditions. Using these “clean” CF zebrafish models, we therefore sought to establish whether a dysfunctional CFTR could regulate non-infectious inflammation *in vivo*.

Neutrophil-mediated inflammation is the characteristic abnormality in the CF airways *(21)*. Therefore, we first investigated the consequence of loss of CFTR on the neutrophilic response throughout a sterile inflammatory process by taking advantage of the reporter transgenic line *TgBAC(mpx:EGFP)i114* with GFP-labelled neutrophils *(18)* (Fig. 1A). Both *cftr* -/- mutant and *cftr* morphant zebrafish larvae displayed overactive neutrophilic responses, typified by early increased and sustained accumulation of neutrophils at wounds, despite the lack of infection (Fig. 1, B and C). While WT fish exhibited a peak number of recruited neutrophils at 4 hours post injury (hpi), neutrophil influx reached a peak at 8 hpi in the absence of CFTR. Since there is evidence of a basal elevation of neutrophil numbers in CF lungs *(4)*, we looked at the global number of neutrophils in *cftr* -/- mutant and *cftr* morphant zebrafish larvae. Although CF animals exhibited a slight increase in the number of neutrophils (Fig. 1, D and E), fMLP-stimulated neutrophil chemotaxis revealed that CF neutrophil responses are indistinguishable from their control counterparts (Fig. 1F) *(13)*. These results suggest that elevation in neutrophil numbers at site of injury is neither due to an overall increase in neutrophils within CF fish nor to a generalised upregulation of chemotaxis in CF neutrophils. Importantly, *cftr* morphants successfully phenocopy the *cftr*-null mutant inflammatory phenotypes, thus validating the use of both CRISPR-Cas9 and morpholino procedures to further investigate the effects of CFTR ablation on host inflammatory responses.

**Fig. 1.**
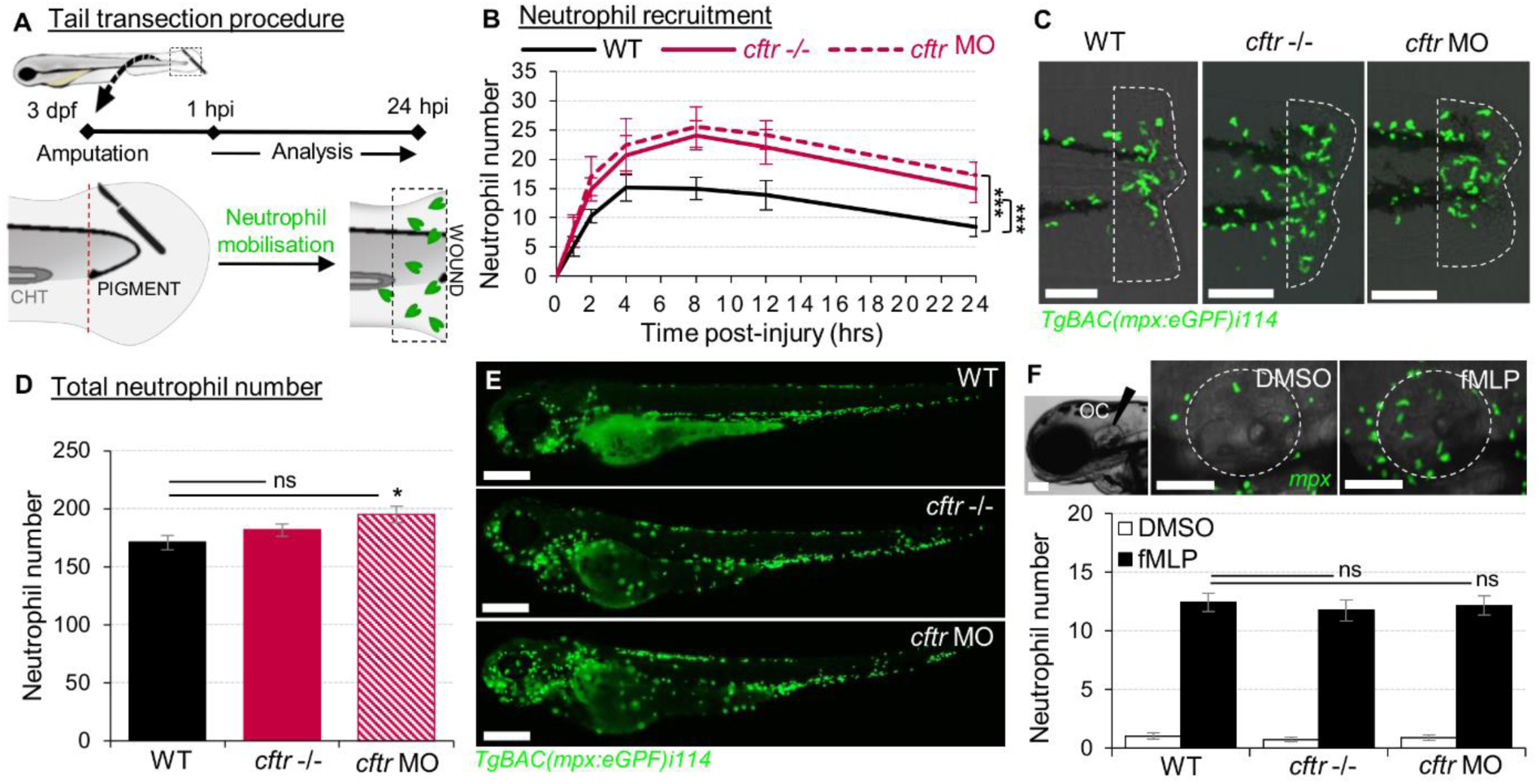
Sterile injury-induced neutrophilic inflammatory responses are exacerbated in the absence of CFTR. (A-C) Neutrophil chemotaxis to sterile tissue lesions in WT, *cftr* -/- mutant (*cftr* -/-) and *cftr* morphant (*cftr* MO) *TgBAC(mpx:EGFP)i114* zebrafish larvae. (A) Larvae were tail amputated then the number of neutrophils mobilized to the site of injury (dotted lines) has been observed and counted by fluorescent microscopy throughout inflammation. The wound area is defined as the region between the amputation edge and caudal hematopoietic tissue end (CHT). (B) Dynamics of neutrophil recruitment towards the wound over 24 hours (n=30; two-way ANOVA with Tukey post-test). (C) Representative confocal images of injured tails at 4 hpi (scale bars, 200 μm). (D-E) Total number of neutrophils (D) in whole *TgBAC(mpx:EGFP)i114* larvae (n=18; one-way ANOVA with Dunnett’s post-test). (E) Microscopy revealed a disorganized neutrophilic distribution in CFTR-depleted animal compared to control counterpart (scale bars, 200 μm). (F) Neutrophil mobilization into the otic cavity (oc, dotted lines) in response to DMSO or fMLP injection in *TgBAC(mpx:EGFP)i114 larvae* monitored at 2 hours-post injection. Representative photomicrographs of fMLP-injected animal compared to DMSO-injected control (top panel; scale bars, 50 μm). Neutrophil counts (n=18; one-way ANOVA with Dunnett’s post-test) (bottom panel).

Collectively, these findings show that sterile injury-induced neutrophilic inflammation is exacerbated in the absence of CFTR, supporting the view that CFTR plays an essential role in control of host innate immunity under normal conditions and revealing a direct link between CF mutations and the inflammatory phenotype in CF in the absence of infection.

### Enhanced ROS production promotes exuberant neutrophil chemotaxis in the absence of *cftr in vivo*

We next asked how CFTR orchestrates neutrophil trafficking during wound-induced inflammation. We first sought to identify the mechanisms driving increased neutrophil infiltration to injury. In CF lungs, high concentrations of chemotactic factors, including IL-8, secreted by epithelial and innate immune cells, are thought to be responsible for the sustained accumulation of neutrophils *(22)*. Despite increased *cxcl8* (interleukin-8) expression following injury in CF fish compared with WT animals (Fig. S2A), neutrophil migration in response to exogenous IL-8 stimuli was similar in WT and CF larvae (Fig. S2B). These results suggest that early changes in neutrophil infiltration in CFTR-deficient zebrafish may be in part altered by cytokines such as IL-8, independently of surface receptor expression such as CXCR1 or CXCR2 in CF neutrophils. However, although IL-8 production was required for amplification and maximal neutrophil accumulation during inflammation, inhibition of IL-8 signaling slightly affected recruitment of early neutrophils towards wounds (Fig. S2C). These findings suggest that CFTR likely regulates the earliest phase of neutrophil mobilisation to injuries in an IL-8-independent manner. We therefore hypothesised that another pro-inflammatory mediator could be abnormally regulated, thus influencing and driving neutrophil inflammation in CF.

Epithelial release of H_2_O_2_ through the respiratory system dual oxidase DUOX2 has been identified as a potent trigger of sterile neutrophilic inflammation *(23, 24)*. Thus, we examined whether loss of CFTR influenced epithelial oxidative responses after tail amputation. While tail transection in WT animals consistently triggered normal oxidative response at injury sites, there was higher ROS production at injured tissues in CFTR-defective fish, demonstrated by increased CellROX staining at wounds (Fig. 2, A and B). Given the importance of DUOX2-derived ROS production in neutrophil chemotaxis, an excessive oxidative response following tissue injury in CF might explain the neutrophilic inflammatory phenotype in CF. qRT-PCR analysis revealed similar *duox2* expression in both injured-WT and CF fish (Fig. 2C). By manipulating oxidative signaling, we next asked whether CFTR/ROS dependent innate immunity is required to orchestrate neutrophilic responses during sterile inflammation. Inhibition of ROS generation by genetic ablation of epithelial-specific DUOX2 oxidase, either though injection of the *duox2*-MO (Fig. 2, D and E) or using diphenyleneiodonium (DPI) (Fig. 2F) *(23)*, strongly reduced early neutrophil chemotaxis to tissue injury in CF animals (Fig. 2G). In contrast, H_2_O_2_-treated WT larvae were characterized by increased numbers of neutrophils at the wound, comparable to that observed in CF fish (Fig. 2F). However, persistence of a neutrophilic response did not appear to be ROS dependent, since blocking the late redox activity with DPI did not reverse wound-associated neutrophil number at later time points in CF zebrafish (Fig. 2H).

**Fig. 2.**
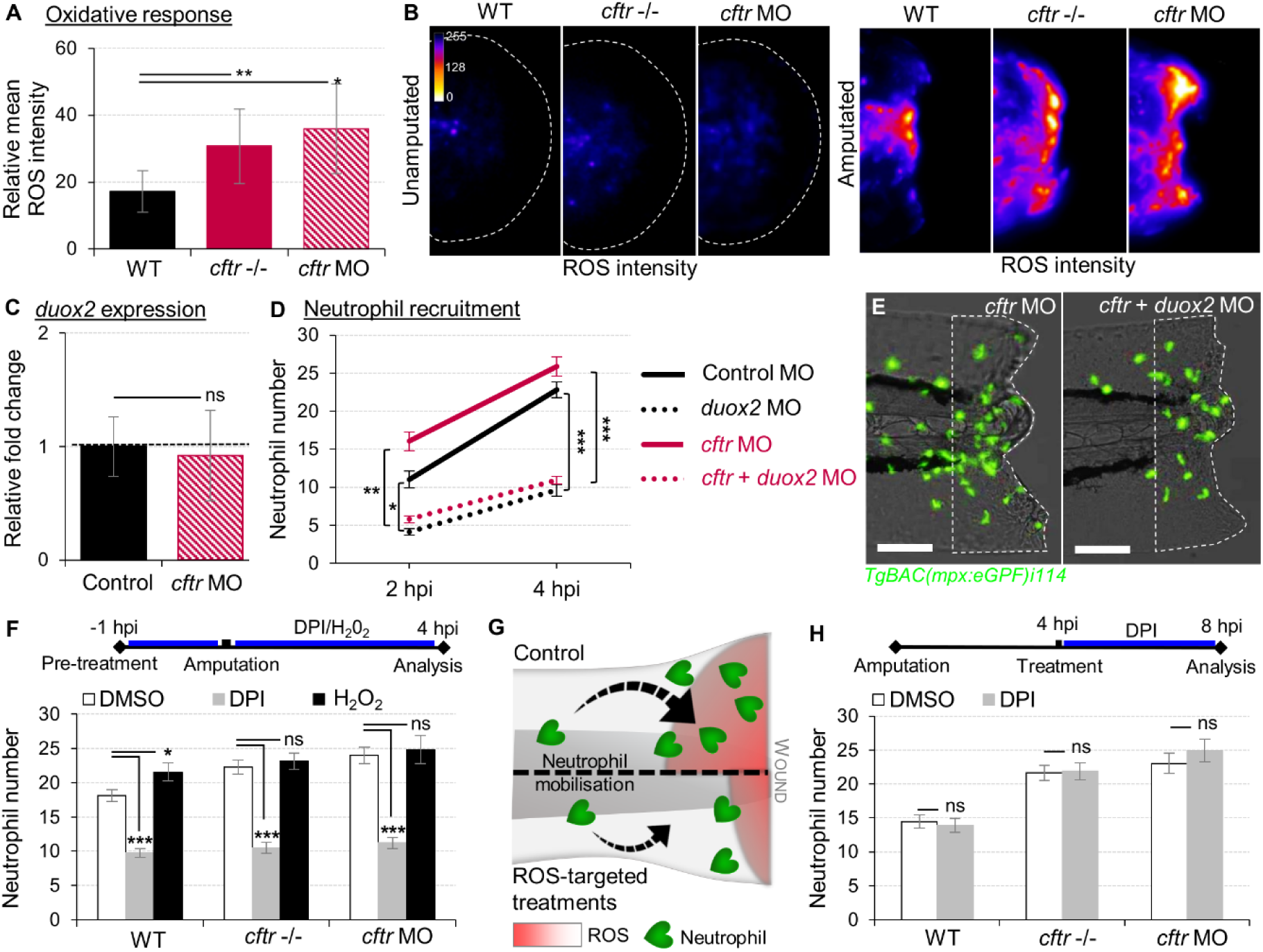
Exuberant wound-induced oxidative responses drive the overactive neutrophilic response in CF larvae. (A-B) WT, *cftr* -/- and *cftr* MO stained with CellROX® to label H_2_O_2_ production. Means ± SEM ROS intensity at 30 minutes post-injury (mpi) (A) and associated pseudocolored photomicrographs of uninjured and injured tails revealing ROS production (B) (n=15; one-way ANOVA with Dunnett’s post-test). (C) mRNA levels of *duox2* gene in tail fin tissue at 30 mpi. Gene expression was expressed as fold change over tail fin tissue from uninjured larvae (30 fins per replicate; mean relative ± SEM gene expression; two-tailed Student t-test). (D) *TgBAC(mpx:EGFP)i114* controls, *cftr, duox2*, and double *cftr/duox2* morphants were tail amputated and neutrophils at wounds were enumerated at 2 and 4 hpi. (n=21; two-way ANOVA with Tukey post-test). (E) Representative photomicrographs of injured tails at 4 hpi (scale bars, 200 μm). (F) WT, *cftr* -/- and *cftr* MO *TgBAC(mpx:EGFP)i114* larvae were pretreated with DPI or H_2_O_2_ prior tail amputation procedure, then injured and immediately put back in treatments. Neutrophil counts at 4 hpi (n=21; one-way ANOVA with Dunnett’s post-test). (G) Schematic diagram showing inhibition of ROS signaling efficiently reduces neutrophil inflammation at wounds in CF animal. (H) *TgBAC(mpx:EGFP)i114* larvae were treated with DPI at 4 hpi. Neutrophil counts at 8 hpi (n=21; two-tailed Bonferroni t-test).

Altogether, these findings indicate that functional CFTR is necessary to orchestrate the early phases of neutrophil recruitment through ROS-mediated mechanisms at sites of damage, and support that deleterious changes in epithelial oxidative responses in CF airway *(25)* are involved in part in the neutrophilic inflammatory phenotype in CF.

### Neutrophil apoptosis and reverse migration are suppressed in CF animals

The resolution of inflammation is brought about by coordinated pro-resolution events acting together to restore tissue homeostasis. These events include: *i)* cessation of neutrophil infiltration and initiation of reverse migration, *ii)* promotion of neutrophil apoptosis, and *iii)* removal of apoptotic inflammatory cells (efferocytosis) *(26)*. In order to elucidate the cellular processes underlying persistent neutrophilic inflammation in CF, we further investigated innate immune cell activity and functions over the time course of inflammation resolution in CFTR-deficient animals.

Firstly, we compared WT and CF animals, to determine any difference in neutrophil apoptosis at injured sites. Combining TUNEL (Fig. 3, A and B) and acridine orange (AO) assays (Fig. S3), confocal analysis revealed that apoptotic neutrophils are found in both the presence and absence of CFTR. However, more importantly, neutrophil apoptosis in CFTR-defective larvae was markedly lower when compared with their control counterpart at 4 and 8 hpi, implying that CFTR plays a crucial role in determining neutrophil lifespan during inflammation.

**Fig. 3.**
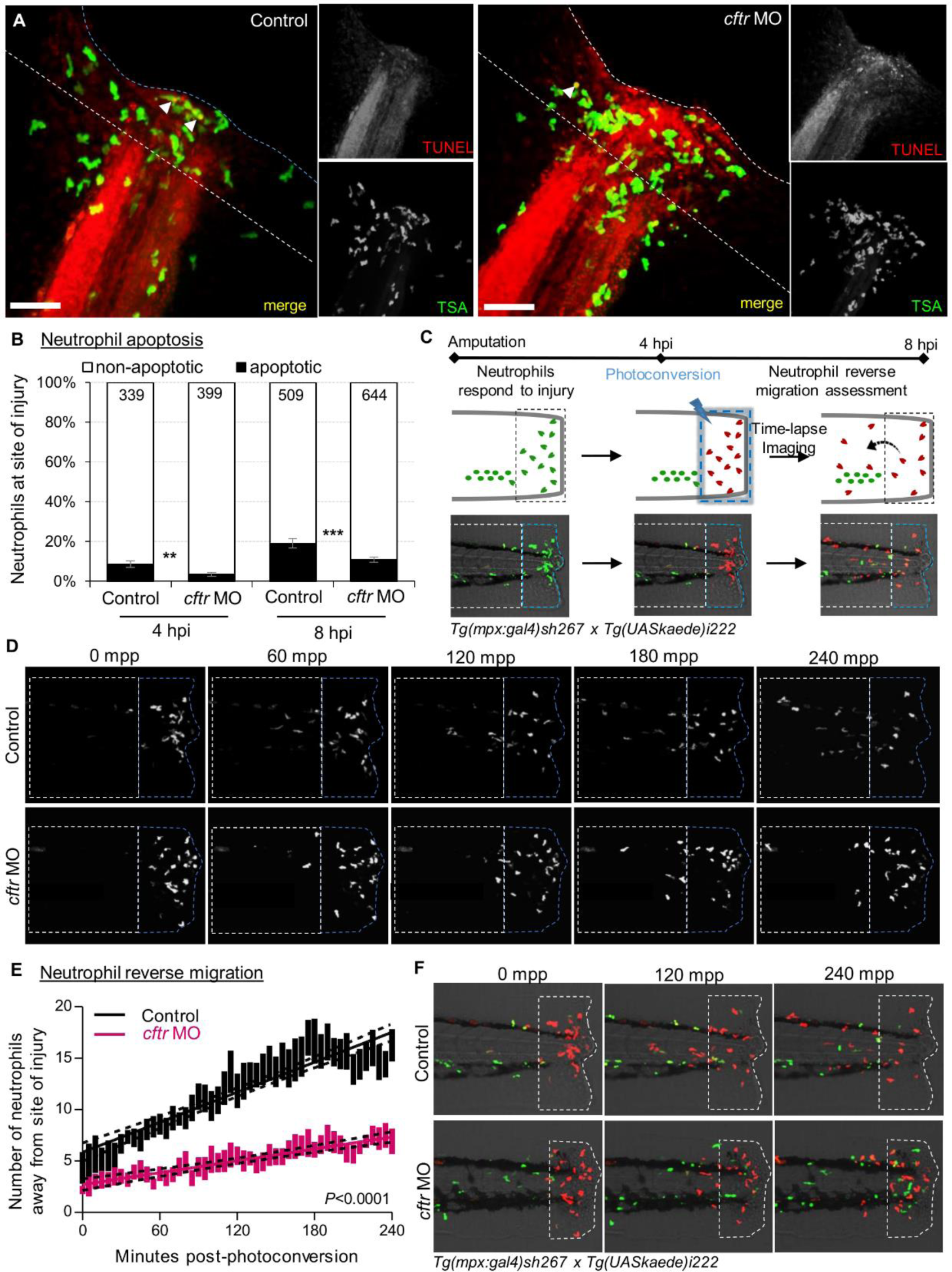
CFTR deficiency delays inflammation resolution *in vivo* both by reducing neutrophil apoptosis and reverse migration of neutrophils. (A-B) *C*ontrol and *cftr* MO larvae were amputated and stained with TUNEL/TSA to label apoptotic cells. (A) Representative confocal pictures of injured tails at 8 hpi (scale bars, 60 μm) revealing the proportion of apoptotic neutrophils at the wound (white arrow). (B) Quantification of neutrophil apoptosis at site of injury at 4 and 8 hpi. (n=18-22, Fisher t test). (C-F) Neutrophil reverse migration in control and *cftr* MO larvae. (C) Tail transection was performed on 3 dpf *Tg(mpx:gal4)sh267;Tg(UASkaede)i222* larvae. The site of injury was photoconverted at 4 hpi, then the number of photoconverted neutrophils (red) moving away (white dotted box) from the wound (blue dotted box) were time-lapse imaged and quantified over 4 hours post photoconversion by confocal microscopy. (D) Representative confocal imaging of amputated tails showing the kinetics of photoconverted neutrophils that migrate away from photoconverted region over inflammation resolution. (E) Plot showing the number of photoconverted neutrophils leaving the area of injury over 4 hours post photoconversion in control and *cftr* MO. Line of best fit shown is calculated by linear regression. *P* value shown is for the difference between the 2 slopes (n=12, performed as 3 independent experiments). (F) Representative confocal imaging of amputated tails showing the kinetics of new neutrophils (green) recruited towards site of injury after photoconversion.

We next addressed if the absence of CFTR affected reverse migration of neutrophils from lesions, as a potential contributor delaying resolution of inflammation in CF. *cftr* was knocked-down in *Tg(mpx:Gal4)sh267;Tg(UAS:Kaede)i222* zebrafish, a neutrophil-specific transgenic line expressing a photoconvertable pigment which can be used to track specific groups of cells. Inflammation in this line was induced by tail amputation, neutrophils at the wound photoconverted at 4 hpi and followed for a further 4 hours. We observed a reduced number of neutrophils migrating away from the injury site in *cftr* morphants (Fig. 3, C and D), demonstrating the importance of functional CFTR for permitting reverse migration and subsequent resolution of inflammation. Additionally, microscopy analysis showed the presence of more late-migrating neutrophils (recruited to the wound after photoconversion), suggesting that defective CFTR activity also delays cessation of neutrophil recruitment (Fig. 3E).

Our recent work has highlighted the crucial role of macrophages in apoptotic cell clearance, for successful resolution of neutrophilic inflammation in zebrafish larvae models *(27)*. Ineffective efferocytosis in CF macrophages has been previously reported *(28)*. To exclude abnormal macrophage behaviour and activity as the cause of delayed resolution of inflammation in CF animals, we therefore investigated macrophage responses to tissue injury using the macrophage-specific *Tg*(*mpeg1*:*mCherry-CAAX*)*sh378* reporter line *(29)*. Firstly, macrophage counts at the site of injury showed similar dynamics of cell recruitment towards wounds over the time course of inflammation in both control or *cftr* morphants, suggesting that loss of CFTR does not affect macrophage mobilization to inflamed tissues (Fig. S4, A and B). We next analysed rates of efferocytosis by macrophages and compared these at 4 and 8 hpi by confocal microscopy, and found no difference in the amount of efferocytosis between *cftr* morphants and controls (Fig. S4, C and D). These results suggest that macrophage activity, which is thought to be important for initiating inflammation resolution, is less dependent on CFTR, suggesting this pathway is not required for the early macrophage responses to a sterile lesion.

Collectively, these findings indicate that loss of CFTR delays resolution of inflammation *in vivo* by reducing neutrophil apoptosis and their reverse migration in the context of sterile tissue inflammation, and thus linking a CFTR-related defect in both neutrophil apoptosis and reverse migration of neutrophils as pathogenic mechanisms leading to persistent neutrophilic inflammation in CF.

### CF-related inflammation impedes successful tissue repair

Resolution of inflammation plays a pivotal role preventing chronic inflammation and excessive tissue damage, as well as initiating tissue healing and repair *(30, 31)*. The profound inflammatory phenotype, prompted us to analyze tissue repair in CF animals. As a developing organism, tissue repair responses in zebrafish larvae are robust, with zebrafish larval tail fins repairing completely over 48-72 hours after amputation, restoring both size and shape (Fig. 4A) *(32)*. We therefore studied the role of CFTR in tissue repair potential and interrogated how unresolved neutrophilic inflammation could be involved in defective tissue repair in CF. By measuring fin areas at 3 dpi, our results showed that CF fish undergo tail fin regrowth by a reduction of 30% (Fig. 4, A and B). Moreover, regenerated tissues in CF animals consistently showed abnormal shape and evidence of damage (Fig. 4B). We hypothesised that harmful neutrophilic activity at sites of injury might contribute to a non-healing wound. Therefore, in order to evaluate the influence of neutrophilic responses on tissue repair, we ablated neutrophils in zebrafish embryos using the *csf3r* morpholino *(33)*. Removal of neutrophils at wounds during inflammation significantly improved tail fin repair in injured *csf3r* morphants in CF (Fig. 4C). Interestingly, reducing early neutrophil infiltration by DPI partially restored tissue regeneration in CFTR-depleted animals (Fig. 4D). Defective tissue repair was not reversed by genetic inhibition of DUOX2 (Fig. 4E). In addition, our results showed that regenerated tail fin area in WT animals is significantly reduced in the presence of exogenous H_2_O_2_ (Fig. 4D), supporting our hypothesis that deleterious neutrophilic inflammation contributes to defective tissue repair in CF.

**Fig. 4.**
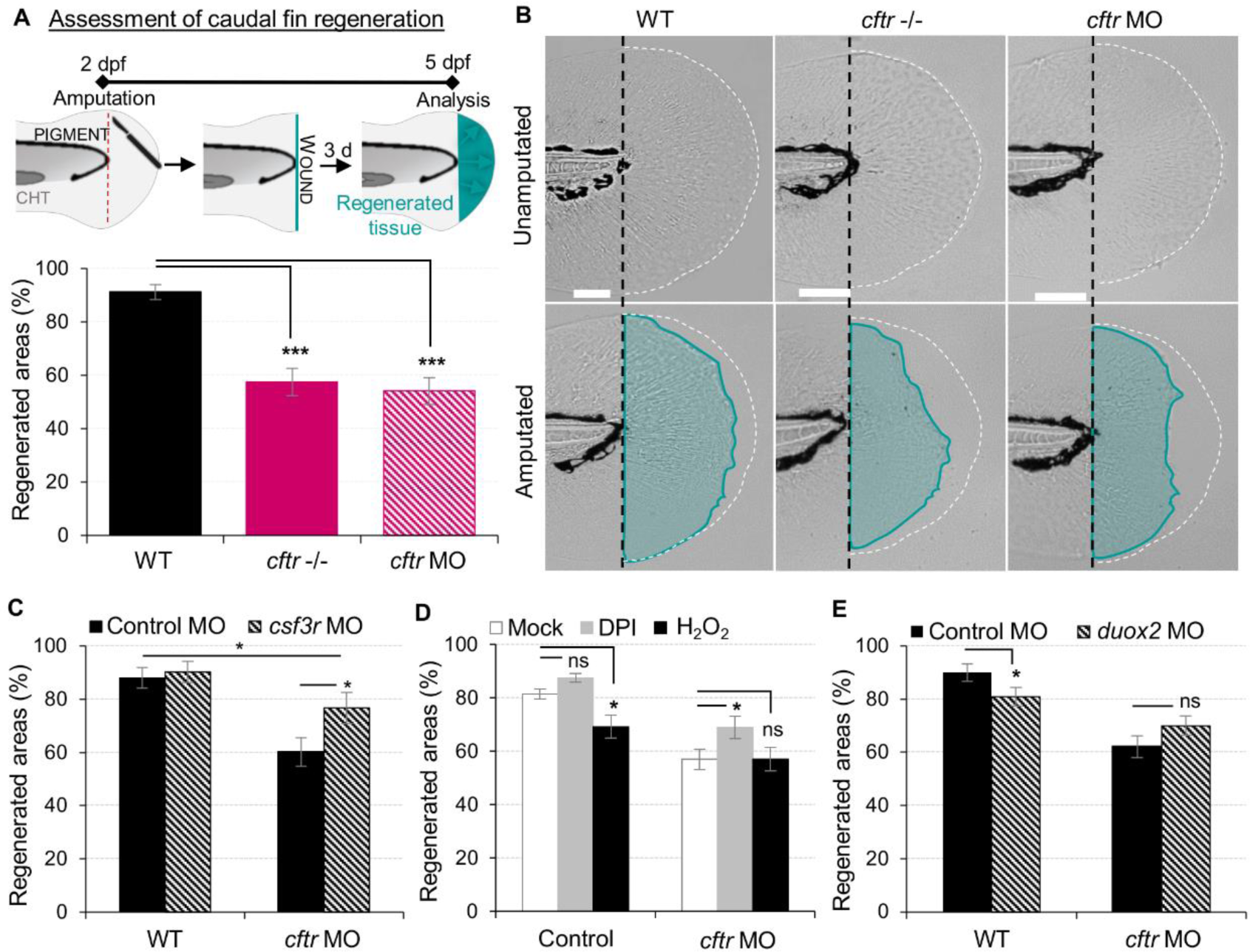
Neutrophilic response in CF hampers tissue repair *in vivo*. (A-B) Tail fin regeneration assessment in WT, *cftr* -/- and *cftr* MO zebrafish. 2 dpf embryos were tail fin amputated then the potential of tail fin regeneration is evaluated by measuring regenerated fin areas (blue). (A) Measurement of regenerated fin areas (n=30, one-way ANOVA with Dunnett’s post-test). (B) Representative imaging of injured tail fin at 3 dpi. (scale bars, 200 μm). (C) Selective ablation of neutrophils has been carried out in WT and CF animals by injecting *csf3r*-MO. Measurement of regenerated fin areas in the presence or absence of neutrophils (n=21, one-way ANOVA with Tukey post-test). (D) *TgBAC(mpx:EGFP)i114* larvae were pretreated with DPI or H_2_O_2_ prior to tail fin amputation procedures, then injured and immediately put back in treatments for 4 hours. Regenerated fin areas are measured at 3 dpi. (n=21, one-way ANOVA with Dunnett’s post-test). (E) Inhibition of NADPH oxidase activity by injecting *duox2*-MO has been carried out in WT and CF animals. Embryos were tail fin injured and regenerated fin areas measured at 3 dpi (n=21, two-tailed Bonferroni t-test).

Collectively, these results indicate that functional CFTR is required for tissue repair, potentially explaining the severe nature of lung destruction in CF compared to other forms of ciliary dysfunction. Additionally, our findings demonstrate that local inflammation mediated by sterile injury and its timely resolution are both crucial for successful tissue repair. These results suggest that the deleterious inflammatory processes impeding tissue performance might be a potential therapeutic target in CF.

### Tanshinone IIA promotes resolution of neutrophilic inflammation and subsequent tissue repair in a CF model

Reducing the impact of inflammation-mediated tissue damage is a major concern in CF therapy. In particular, therapeutic targeting of inflammation resolution is an attractive strategy to bring about healing and tissue repair, particularly if we can do so without compromising host defences against pathogens. We have previously identified the potential of Tanshinone IIA (TIIA) to accelerate resolution of inflammation by enhancing both reverse migration and apoptosis of neutrophils *(34)*. We therefore investigated whether TIIA treatment could resolve wound-induced inflammation and initiate regenerative responses in a CF context. Interestingly, while some reports have highlighted the anti-oxidative properties of TIIA *(35)*, our results showed that early neutrophil mobilization towards wounds was not affected by TIIA exposure in CF zebrafish (Fig. 5A), and are reminiscent of our results in WT fish *(34)*. However, when TIIA was used during the resolution phase of inflammation, TIIA treatment strongly reduced wound-associated inflammation in CF animals, as determined by neutrophil numbers (Fig. 5, B and C). In order to further understand the mechanism of inflammation resolution mediated by TIIA in a CFTR-deficient context, we next studied neutrophil activity during inflammation resolution. Our results showed that TIIA enhanced both neutrophil apoptosis (Fig. 5D) and migration of neutrophils away from wounds (Fig. 5E) in injured CF larvae. Remarkably, CFTR-deficient larvae treated with TIIA exhibited markedly enhanced tissue repair of the tail fin (Fig. 5, F and G).

**Fig. 5.**
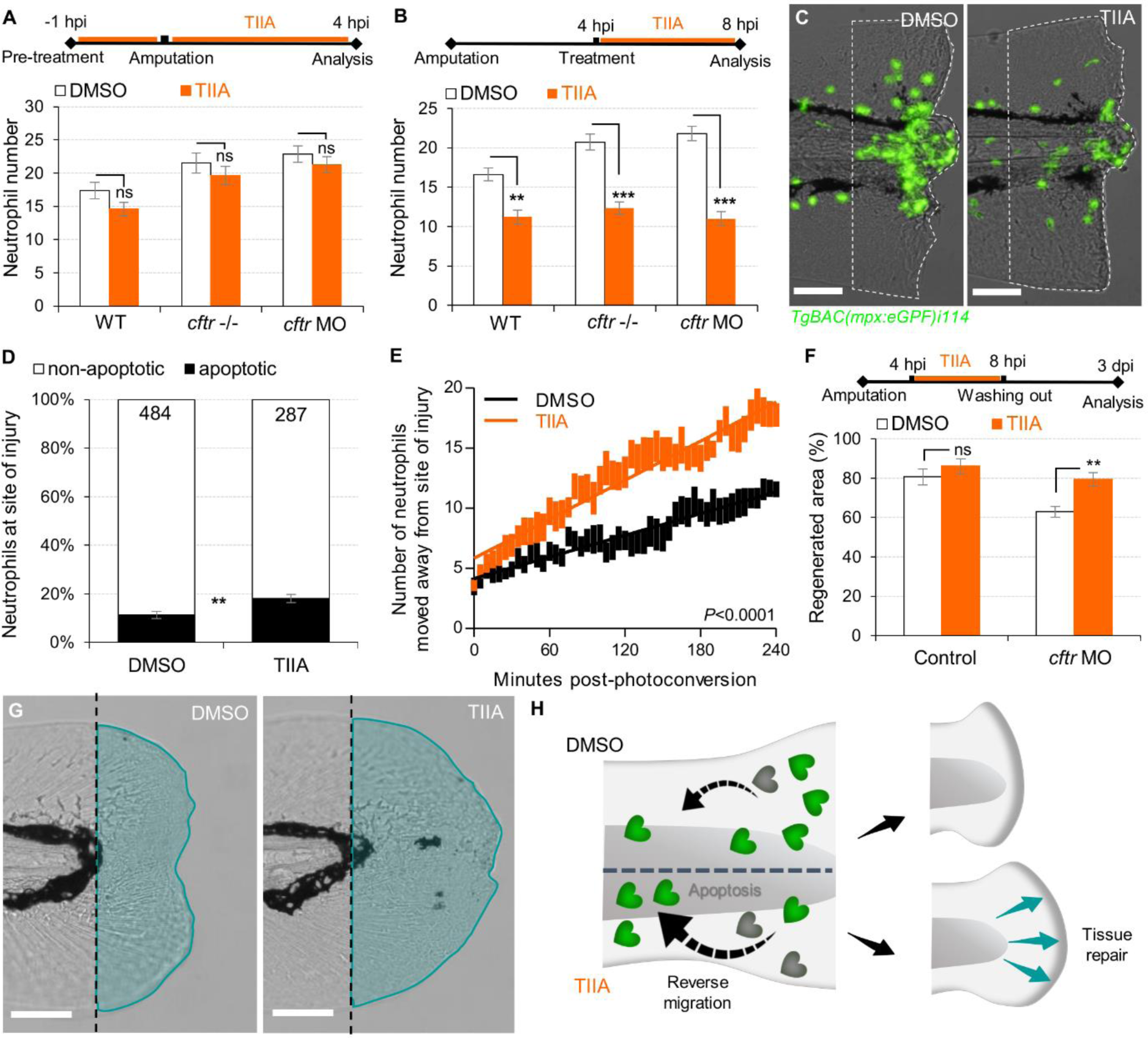
TIIA-driven neutrophil apoptosis and reverse migration accelerate inflammation resolution in CF. (A) *TgBAC(mpx:EGFP)i114* larvae were pretreated with 25µM of TIIA prior to tail fin amputation procedure, then injured and immediately put back in treatments for 4 hours. Neutrophil number at the wound was counted at 8 hpi (n=21, two-tailed Bonferroni t-test). (B-C) *TgBAC(mpx:EGFP)i114* larvae were injured and treated from 4 hpi with 25µM of TIIA. (B) Neutrophil number at wound was counted at 8 hpi (n=21, two-tailed Bonferroni t-test). (C) representative number of neutrophils remaining at wounds at 8 hpi (scale bars, 200 μm). (D) Neutrophil apoptosis quantification at 8 hpi in *cftr MO* treated with 25 µM of TIIA from 4 hpi and stained with TUNEL/TSA. (n=40, Fisher t test) (E) Reverse-migration assay in *cftr* MO *Tg(mpx:gal4)sh267;Tg(UASkaede)i222*. At 4 hpi fish were treated with 25 µM of TIIA and neutrophils at site of injury were photoconverted. The numbers of photoconverted cells that moved away from the wound were time-lapse imaged and quantified over 4 hours. (F-G) Regenerative performance after TIIA treatment. (F) Regenerated fin areas are measured at 3 dpi. (n=21, two-tailed Bonferroni t-test). (G) Representative imaging of injured tail fin at 3 dpi. (scale bars, 200 μm). (H) Schematic diagram showing TIIA efficiently accelerates inflammation resolution by inducing neutrophil apoptosis and reverse migration at wounds and improves tissue repair in CF animal.

Altogether, these data demonstrate that redirecting neutrophils to apoptosis and reserve migration using TIIA may be a targeted therapeutic strategy to restore tissue repair (Fig. 5H) and thus prevent inflammatory lung damage in CF.

## Discussion

Despite considerable improvement in the care of inflammatory pathology in CF patients, inflammation-related progressive pulmonary destruction leading to respiratory failure remains the leading causes of premature death in CF. Our understanding of the mechanistic links between CFTR mutations and the pathogenesis of inflammatory pulmonary disease in CF is far from complete. The *in vivo* evaluation of biological functions of CFTR in currently available CF models suffers from several limitations, predominantly the evaluation of phenomena in a pre-existing inflammatory environment (including epigenetic scars). To overcome this, we established and exploited a transparent zebrafish larval model, as a physiologically-relevant sterile tissue damage model *in vivo*, where innate immune responses can be readily imaged and manipulated. Using CFTR-depleted zebrafish, we have investigated the effects of CFTR dysfunction on host innate immune response to tissue injury, and uncovered a number of altered inflammatory processes as critical mechanisms underlying inflammatory disease in CF.

The spatiotemporal events associated with CFTR ablation reveal a mechanism whereby CFTR participates in the adjustment of innate immunity, conditioning key regulatory mediators involved in the regulation of inflammation and tissue repair (Fig. 6). Even in the absence of invading pathogens, the innate immune response is initially triggered by danger signals released from damaged cells. This study suggests that inflammation, irrespective of the initiating stimulus is exacerbated in the absence of CFTR. This is in marked contrast to the classical explanations linking excessive inflammation to chronic infection, providing a new potential explanation for how CFTR directly regulates host inflammatory potential *in vivo*. This observation suggests that targeting ROS and neutrophils directly might have a much larger effect on inflammatory tissue damage than previously supposed. Without neutrophilic inflammation to generate increased extracellular DNA release from the dead pulmonary cells, this might result in major improvements in mucus viscosity and hence pathogen handling in CF.

**Fig. 6.**
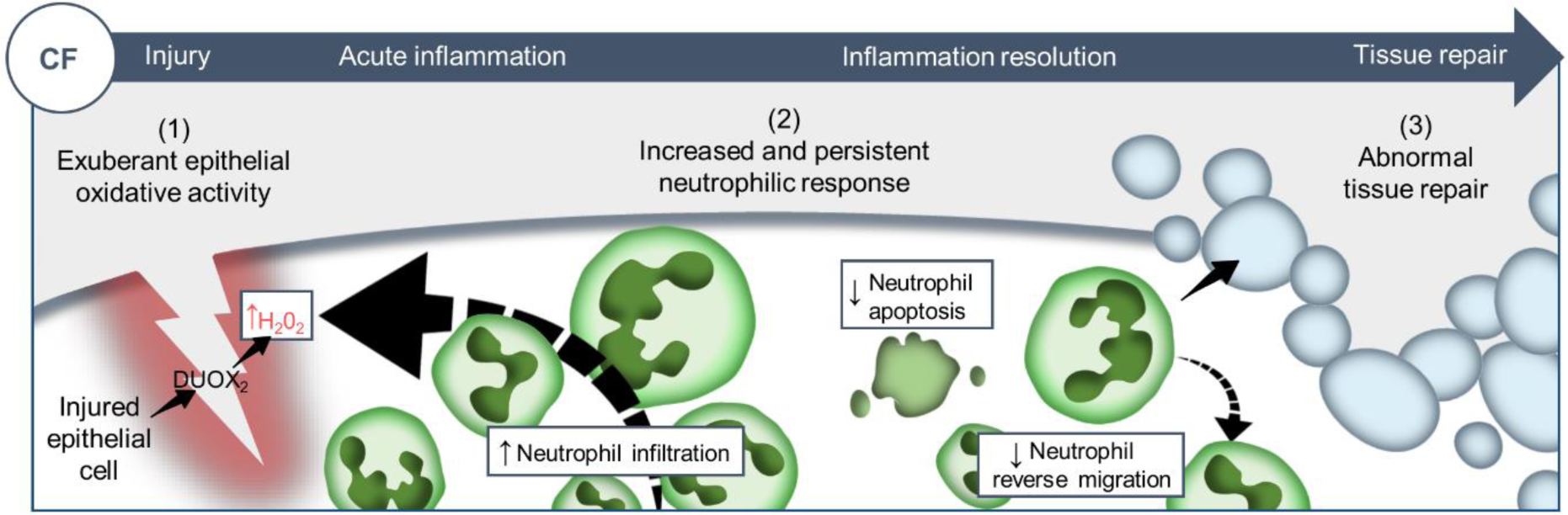
Proposed model showing CF-related inflammation immuno-pathogenesis. A sterile lesion is characterized by early release of epithelial H_2_O_2_ through DUOX2, leading to neutrophil recruitment towards wounds; coordination of neutrophil apoptosis and reverse migration promoting efficient inflammation resolution, and allowing to restore tissue homeostasis and initiate tissue repair. In contrast, in CF zebrafish sterile injury leads to extensive inflammation, typified by increased then sustained accumulation of neutrophils at wounds: (1) excessive epithelial ROS release drives increased neutrophil recruitment towards wounds; (2) reduction of neutrophil apoptosis and impaired retrograde migration of neutrophils resulting in delayed resolution of inflammation. (3) Therefore, the increased number of neutrophils that mobilize in an uncontrolled manner at wound sites causes persistent inflammation, severe tissue damage, and abnormal tissue repair.

H_2_O_2_ produced in epithelia is a potent chemoattractant source for neutrophils *(23, 36)* and has been proposed as an important component in the neutrophilic inflammatory pathogenesis of CF lung disease *(25)*. Our work highlights that loss of CFTR leads to an overactive neutrophilic response to tissue damage, mediated by excessive epithelial ROS generation, and demonstrates that CFTR/DUOX2 axis-dependent ROS production is instrumental in efficiently orchestrating neutrophil chemotaxis (Fig. 2). Surprisingly, such exuberant neutrophil mobilisation is not observed when CFTR-defective zebrafish are infected with CF pathogens *(12, 13)*, strongly implying tissue damage as the primary initiator in increasing inflammation rather than infection in CF. This observation is of particular interest in light of clinical reports of human patients showing early elevation of neutrophil infiltration in CF airway in the absence of detectable infection *(4)*. Whether CFTR/DUOX2 NADPH axis differentially regulates sterile or infection-induced neutrophilic inflammation remains to be addressed.

While progress has been made, the mechanistic link between dysfunctional CFTR and abnormal ROS generation is still elusive with evidence for both negative and positive regulation depending on the cell type. For example, our recent study shows that CF zebrafish exhibit increased susceptibility to *M. abscessus* in part due to an inability to generate effective oxidative immunity in professional phagocytes *(13)*, thus indicating opposite effects on oxidative responses in myeloid cells and in epithelial cells as shown in this present work. Further investigation is warranted to find the as yet unidentified molecular basis and to delineate the differential role of the CFTR/ROS axis in both epithelial inflammation and immunity to infection, and the link with CF phenotype in patients.

Mechanisms leading to efficient inflammation resolution, tissue healing and repair depend on suppression of inflammatory signaling pathways, orchestrated by a tightly regulated innate immune response. CF lung disease is characterised by an unresolved inflammatory response. Corroborating data from other CF *ex vivo* models *(21, 37)*, we show that CF neutrophils that have migrated towards inflamed sites display molecular changes associated with reductions in both apoptosis and reverse migration, and thus have enhanced potential to drive tissue damage and impair tissue repair because of their prolonged survival and extended activity at the wound. Although it is unclear whether these are primary neutrophil defects or a response to wound-mediated inflammation in CF zebrafish, these results emphasize that CFTR plays an important role in the maintenance of tissue homeostasis by determining neutrophil behaviour and lifespan during inflammation process.

Tissue repair after injury depends on both host regenerative capacity and the quality of the inflammatory response *(30, 31)*. Reminiscent of previous studies suggesting that CFTR plays a critical role in wound repair *(38, 39)*, we show here that CF zebrafish exhibit incomplete tissue repair after tail-fin amputation, which can be improved by genetic ablation of neutrophils. These results indicate the deleterious role of neutrophil activity contributing to tissue damage and impaired tissue repair in the context of CF. Importantly, this also appears to be ROS dependent, since therapeutic approaches modulating oxidative responses markedly reduce neutrophilic inflammation to tissue injury *in vivo* as well as greatly decrease collateral damage from inflammation and improve tissue repair. However, if oxidative stress is considered a causal mediator of damage and inflammatory disease, ROS are crucial regulators of regenerative inflammation *(40, 41)*, as suggested by genetic inhibition of the DUOX2 NADPH oxidase pathway modulating neutrophil response at wounds but interfering with tissue repair processes. These findings emphasize that balancing the positive and negative effects of the inflammatory process should be considered for the design of clinical treatments for the management of inflammatory disease in CF. Importantly, while tissue repair in CF zebrafish can be improved by re-balancing neutrophilic activity at wounds, their regenerated tissue area remains smaller than in CF controls. This suggests that CF-mediated alterations other than unresolved inflammation are responsible for the impaired tissue repair in CF fish and thus indicates that additional CFTR-mediated mechanisms are likely to participate in host tissue repair capacity. The molecular basis of defective tissue repair in CF is particularly intriguing and deserves further attention, and undoubtedly will be crucial to support optimal tissue repair in CF.

Controlling injurious effects of inflammation is an essential component in the management of CF. Overall, our findings indicating that CF neutrophils have prolonged survival and activity due to dysfunctional apoptosis *(37, 42)* and reverse migration, providing an explanation as to how CFTR mutations may lead to unresolved neutrophilic responses, impaired tissue repair resulting in scar formation or fibrosis in CF lung, and highlighting various approaches that aim at modulating these mechanisms to limit inflammation-driven tissue damage and promote tissue repair. Assuming that this dysregulated innate immunity, prior to bacterial colonisation, disrupts both inflammatory responses and tissue repair in CF, therapeutic strategies to normalise harmful neutrophilic inflammation might simultaneously promote resolution of inflammation and tissue repair, and thus prevent pulmonary destruction. Current strategies based on anti-inflammatory treatments in CF, including ibuprofen or corticosteroids, have not yet proven effective in the clinic and also carry persistent long-term use safety concerns *(43)*. TIIA, derived from the traditional Chinese medicinal herb *Salvia miltiorrhiza* widely used for the treatment of patients with cancers, inflammatory or cardiovascular diseases *(44–46)*, was also found to be effective as a pro-resolution compound by inducing apoptosis of neutrophils and promoting their reverse migration in both zebrafish larval model and human neutrophils *(34)*. Considering evidence regarding the dysfunctional responses to apoptosis as a driver of persistent neutrophilic inflammation in the CF lung, we proposed here to study efficiency of TIIA on inflammation outcomes in a context of CF, attempting to restore dysregulated inflammation and epithelial integrity in CF. We show that TIIA can effectively rebalance neutrophilic inflammation in CF animal by counteracting signaling pathways associated with neutrophil persistence and survival. Consequently, TIIA can efficiently prevent inflammatory tissue damages and improve tissue repair. These findings have significant therapeutic implications for potently targeting neutrophilic inflammation in CF, while minimising risk of blocking host immunity, and thus may support existing therapeutic strategies or could be an alternative to existing anti-inflammatory approaches. TIIA is currently used in clinical trials in other respiratory diseases *(47)*, and its effects are conserved in human neutrophils *(34)*, suggesting it might be possible to directly test this hypothesis.

In conclusion, we report here a direct stepwise dissection of the inflammatory response in an animal lacking CFTR, providing a more comprehensive delineation of the cellular basis linking CFTR deficiency with inflammatory pathogenesis of the CF airways, and consequently insights for development of specific therapies aimed at restoring innate immune potential of CF patients and thus identify novel treatment approaches to alleviate neutrophil inflammation-driven tissue damage, with improvement in both quality of life and life expectancy.

## Materials and methods

### Zebrafish Husbandry and Ethic statements

Experimental procedures were performed using the following zebrafish transgenic lines: *TgBAC(mpx*:*eGFP)i114* and *Tg(Lyz:DSred)nz5* labelling neutrophils *(18, 48)*; *Tg*(*mpeg1*:*mCherry-CAAX*)*sh378* labelling macrophages *(29)*; *Tg(mpx:gal4)sh267;Tg(UAS:kaede)i222* zebrafish were used for reverse migration assays *(49, 50)*. To study gene expression, oxidative activity and tissue repair potential, the WT pigment-less *nacre (51)* larvae was used.

Fish were maintained in buffered reverse osmotic water systems at 28°C and were exposed on a 14:10-hour light/dark cycle to maintain proper circadian conditions in UK Home Office-approved facilities in the Bateson Centre aquaria at the University of Sheffield, under AWERB (Animal Welfare and Ethical Review Bodies). All zebrafish experiments described in the present study were performed on larvae <5 dpf, to standards set by the UK Home Office. Maintenance of adult mutant fish was approved under home office license P1A4A7A5E.

### Generation of a stable *cftr* null mutant

Purified gRNA, Cas9 protein, and tracrRNA were purchased from Sigma-Aldrich. We used the following pair of gRNA to generate the *cftr* mutant^sh540^ reported here (PAM site is indicated in bold): **CCC**TCCATCGCGTCTCAGTAGAT and AATCGTCAACCCTCTTGGGG**TGG**. 1nl of gRNA pair was co-injected with Cas9 protein and tracrRNA into the yolk of *TgBAC(mpx*:*eGFP)i114 or* nacre larvae at one-cell-stage as described earlier *(27)*. Genomic DNA was extracted and prepared from individual larvae at 2 dpf as previously described *(53)*. PCR using Firepol® (Solis BioDyne) was used to amplify a 320 bp region into the *cftr* gene (ENSDARG00000041107). Primer sequences used were as follows: *cftr_fw* CCTTTCCTGAGCTTCAGTCAG *cftr_rev* CACCAGGGAGAACTTTCTGTC. WT Mutant forms produce 193-bp band (Fig. S1A). Each founder (F0) is outcrossed with a WT fish, and then embryos from each outcrossed pair (F1 offspring) are screened for germline transmission mutations. Genomic DNA was isolated from 72 hpf tail biopsies *(53)* and heterozygotes mutants were identified *via* gel based genotyping targeted deletion in the *cftr* gene. Heterozygous larvae carrying mutations were used to achieve a stable population of *cftr* +/- adults. *cftr*-/- embryos were produced by crossing these heterozygous lines and screened for impaired Kupffer’s vesicle inflation, as initially reported by Navis et al in a TALEN-generated *cftr-*null mutant *(54)*.

### *In vivo* sterile inflammation assays

All sterile injury-induced inflammation was performed following established methods *(18)*. Briefly, 3 dpf larvae were tricaine-anesthetized, then transection of the tail was performed with a microscalpel (5 mm depth; World Precision Instruments). Neutrophil or macrophage chemotaxis were evaluated by assessing the number of cells at wound sites (the region posterior to the circulatory loop, Fig. 1A) at various time points throughout inflammatory process on a fluorescence dissecting stereomicroscope (Leica).

### Pharmacological treatments for anti-inflammation and pro-resolution assays

Larvae were incubated in sterile E3 media supplemented with 25 μM Tanshinone IIA (TIIA, Sigma-Aldrich) *(34)*, 100 μM Diphenyleneiodonium (DPI, Sigma-Aldrich) *(23)*, H_2_O_2_ (ThermoFisher) *(55)* or 1% Dimethyl sulfoxide (DMSO, Sigma-Aldrich) as negative control. For anti-inflammation procedures, larvae were pretreated with indicated compounds prior tail fin amputation challenge, then injured and immediately put back in treatments. Neutrophils at wounds were counted at 4 hpi at the peak of recruitment. For resolution of inflammation assays, injured larvae were raised to 4 hpi then incubated in E3 media supplemented with indicated compounds. Neutrophils at the wound sites were enumerated at 8 hpi for inflammation resolution.

### *In vivo* tissue repair assay

For regeneration assays, 2 dpf embryos were anesthetized and tail fins were transected at the region indicated in Fig. 4A. Tissue repair performances were evaluated by assessing the regenerated tail fin area at 3 dpi. Percentage of regeneration was calculated by normalizing the regenerated tail fin area versus fin areas of unamputated animal (WT or CF fish).

### Quantification and Statistical Analysis

Statistical analysis was performed using R 3.5.0 (R core team) or Prism 7.0 (GraphPad Software, CA, USA) and detailed in each Figure legend. All data are presented as means from 3 independent experiments. All error bars indicate standard errors of means (SEM). ns, not significant (p≥ 0.05); *p< 0.05; **p< 0.01; ***p<0.001.

## Acknowledgments

We gratefully acknowledge the Bateson Centre aquarium and Wolfson Light Microscopy Facility (MRC grant G0700091 and Wellcome Trust grant (GR077544AIA) at the University of Sheffield. We also would like to thank David Drew for technical assistance and support.

## Author contributions

A.B., R.A.F. and S.A.R. designed the study and analyzed the data. A.B. performed the experiments. C.A.L. provided training, maintenance and breeding of zebrafish lines. A.B., R.A.F. and S.A.R. wrote the manuscript.

## Data and materials availability

All data needed to evaluate the conclusions in the paper are present in the paper and/or the Supplementary Materials. Additional data related to this paper may be requested from the authors. Materials could be obtained through a material transfer agreement.

## References and notes

1. J. M. Rommens, M. C. Iannuzzi, B. S. Kerem, M. L. Drumm, G. Melmer, M. Dean, R. Rozmahel, J. L. Cole, D. Kennedy, N. Hidaka, M. Zsiga, M. Buchwald, J. R. Riordan, L. C. Tsui, F. S. Collins, Identification of the cystic fibrosis gene: Chromosome walking and jumping, Science (80-.). (1989), doi:10.1126/science.2772657.

2. D. C. Gadsby, P. Vergani, L. Csanády, The ABC protein turned chloride channel whose failure causes cystic fibrosis Nature (2006), doi:10.1038/nature04712.

3. S. H. Donaldson, R. C. Boucher, in Principles of Molecular Medicine, (2006).

4. C. Verhaeghe, K. Delbecque, L. de Leval, C. Oury, V. Bours, Early inflammation in the airways of a cystic fibrosis foetus, J. Cyst. Fibros. (2007), doi:10.1016/j.jcf.2006.12.001.

5. N. Aldallal, E. E. McNaughton, L. J. Manzel, A. M. Richards, J. Zabner, T. W. Ferkol, D. C. Look, Inflammatory response in airway epithelial cells isolated from patients with cystic fibrosis, Am. J. Respir. Crit. Care Med. (2002), doi:10.1164/rccm.200206-627OC.

6. G. M. Lavelle, M. M. White, N. Browne, N. G. McElvaney, E. P. Reeves, Animal Models of Cystic Fibrosis Pathology: Phenotypic Parallels and Divergences, Biomed Res. Int. (2016), doi:10.1155/2016/5258727.

7. S. A. Renshaw, N. S. Trede, A model 450 million years in the making: zebrafish and vertebrate immunity, Dis. Model. Mech. (2012), doi:10.1242/dmm.007138.

8. S. A. Renshaw, C. A. Loynes, S. Elworthy, P. W. Ingham, M. K. B. Whyte, in Experimental Lung Research, (2007).

9. J. S. Martin, S. A. Renshaw, Using in vivo zebrafish models to understand the biochemical basis of neutrophilic respiratory disease, Biochem. Soc. Trans. (2009), doi:10.1042/bst0370830.

10. Z. Zhang, J. Chen, Atomic Structure of the Cystic Fibrosis Transmembrane Conductance Regulator, Cell (2016), doi:10.1016/j.cell.2016.11.014.

11. F. Liu, Z. Zhang, L. Csanády, D. C. Gadsby, J. Chen, Molecular Structure of the Human CFTR Ion Channel, Cell (2017), doi:10.1016/j.cell.2017.02.024.

12. R. T. Phennicie, M. J. Sullivan, J. T. Singer, J. A. Yoder, C. H. Kim, Specific resistance to Pseudomonas aeruginosa infection in zebrafish is mediated by the cystic fibrosis transmembrane conductance regulator, Infect. Immun. (2010), doi:10.1128/IAI.00302-10.

13. A. Bernut, C. Dupont, N. V. Ogryzko, A. Neyret, J. L. Herrmann, R. A. Floto, S. A. Renshaw, L. Kremer, CFTR Protects against Mycobacterium abscessus Infection by Fine-Tuning Host Oxidative Defenses, Cell Rep. (2019), doi:10.1016/j.celrep.2019.01.071.

14. M. Bagnat, A. Navis, S. Herbstreith, K. Brand-Arzamendi, S. Curado, S. Gabriel, K. Mostov, J. Huisken, D. Y. R. Stainier, Cse1l is a negative regulator of CFTR-dependent fluid secretion, Curr. Biol. (2010), doi:10.1016/j.cub.2010.09.012.

15. Y. J. Guh, C. H. Lin, P. P. Hwang, Osmoregulation in zebrafish: Ion transport mechanisms and functional regulation, EXCLI J. (2015), doi:10.17179/excli2015-246.

16. A. Navis, M. Bagnat, Loss of cftr function leads to pancreatic destruction in larval zebrafish, Dev. Biol. (2015), doi:10.1016/j.ydbio.2014.12.034.

17. H. Sun, Y. Wang, J. Zhang, Y. Chen, Y. Liu, Z. Lin, M. Liu, K. Sheng, H. Liao, K. S. Tsang, X. Zhang, X. Jiang, W. Xu, M. Mao, H. C. Chan, CFTR mutation enhances Dishevelled degradation and results in impairment of Wnt-dependent hematopoiesis article, Cell Death Dis. (2018), doi:10.1038/s41419-018-0311-9.

18. S. A. Renshaw, C. A. Loynes, D. M. I. Trushell, S. Elworthy, P. W. Ingham, M. K. B. Whyte, Atransgenic zebrafish model of neutrophilic inflammation, Blood (2006), doi:10.1182/blood-2006-05-024075.

19. J. R. Mathias, M. E. Dodd, K. B. Walters, S. K. Yoo, E. A. Ranheim, A. Huttenlocher, Characterization of zebrafish larval inflammatory macrophages, Dev. Comp. Immunol. (2009), doi:10.1016/j.dci.2009.07.003.

20. L. Li, B. Yan, Y. Q. Shi, W. Q. Zhang, Z. L. Wen, Live imaging reveals differing roles of macrophages and neutrophils during zebrafish tail fin regeneration, J. Biol. Chem. (2012), doi:10.1074/jbc.M112.349126.

21. D. G. Downey, S. C. Bell, J. S. Elborn, Neutrophils in cystic fibrosis Thorax (2009), doi:10.1136/thx.2007.082388.

22. T. P. Dean, Y. Dai, J. K. Shute, M. K. Church, J. O. Warner, Interleukin-8 concentrations are elevated in bronchoalveolar lavage, sputum, and sera of children with cystic fibrosis, Pediatr. Res. (1993), doi:10.1203/00006450-199308000-00010.

23. P. Niethammer, C. Grabher, A. T. Look, T. J. Mitchison, A tissue-scale gradient of hydrogen peroxide mediates rapid wound detection in zebrafish, Nature (2009), doi:10.1038/nature08119.

24. I. A. Gamaley, I. V. Klyubin, in (1999).

25. F. Galli, A. Battistoni, R. Gambari, A. Pompella, A. Bragonzi, F. Pilolli, L. Iuliano, M. Piroddi, M. C. Dechecchi, G. Cabrini, Oxidative stress and antioxidant therapy in cystic fibrosis Biochim. Biophys. Acta - Mol. Basis Dis. (2012), doi:10.1016/j.bbadis.2011.12.012.

26. A. Ortega-Gómez, M. Perretti, O. Soehnlein, Resolution of inflammation: An integrated view EMBO Mol. Med. (2013), doi:10.1002/emmm.201202382.

27. C. A. Loynes, J. A. Lee, A. L. Robertson, M. J. G. Steel, F. Ellett, Y. Feng, B. D. Levy, M. K. B. Whyte, S. A. Renshaw, PGE2 production at sites of tissue injury promotes an anti-inflammatory neutrophil phenotype and determines the outcome of inflammation resolution in vivo, Sci. Adv. (2018), doi:10.1126/sciadv.aar8320.

28. R. W. Vandivier, T. R. Richens, S. A. Horstmann, A. M. deCathelineau, M. Ghosh, S. D. Reynolds, Y.-Q. Xiao, D. W. Riches, J. Plumb, E. Vachon, G. P. Downey, P. M. Henson, Dysfunctional cystic fibrosis transmembrane conductance regulator inhibits phagocytosis of apoptotic cells with proinflammatory consequences, Am. J. Physiol. Cell. Mol. Physiol. (2009), doi:10.1152/ajplung.00030.2009.

29. A. Bojarczuk, K. A. Miller, R. Hotham, A. Lewis, N. V. Ogryzko, A. A. Kamuyango, H. Frost, R. H. Gibson, E. Stillman, R. C. May, S. A. Renshaw, S. A. Johnston, Cryptococcus neoformans Intracellular Proliferation and Capsule Size Determines Early Macrophage Control of Infection, Sci. Rep. (2016), doi:10.1038/srep21489.

30. M. Karin, H. Clevers, Reparative inflammation takes charge of tissue regeneration Nature (2016), doi:10.1038/nature17039.

31. A. L. Mescher, A. W. Neff, M. W. King, Inflammation and immunity in organ regeneration, Dev. Comp. Immunol. (2017), doi:10.1016/j.dci.2016.02.015.

32. M. K. Iovine, Conserved mechanisms regulate outgrowth in zebrafish fins Nat. Chem. Biol. (2007), doi:10.1038/nchembio.2007.36.

33. N. Palha, F. Guivel-Benhassine, V. Briolat, G. Lutfalla, M. Sourisseau, F. Ellett, C. H. Wang, G. J. Lieschke, P. Herbomel, O. Schwartz, J. P. Levraud, Real-Time Whole-Body Visualization of Chikungunya Virus Infection and Host Interferon Response in Zebrafish, PLoS Pathog. (2013), doi:10.1371/journal.ppat.1003619.

34. A. L. Robertson, G. R. Holmes, A. N. Bojarczuk, J. Burgon, C. A. Loynes, M. Chimen, A. K. Sawtell, B. Hamza, J. Willson, S. R. Walmsley, S. R. Anderson, M. C. Coles, S. N. Farrow, Y. Solari, S. Jones, L. R. Prince, D. Irimia, G. Ed Rainger, V. Kadirkamanathan, M. K. B. Whyte, S. A. Renshaw, A zebrafish compound screen reveals modulation of neutrophil reverse migration as an anti-inflammatory mechanism, Sci. Transl. Med. (2014), doi:10.1126/scitranslmed.3007672.

35. J. Fu, H. Huang, J. Liu, R. Pi, J. Chen, P. Liu, Tanshinone IIA protects cardiac myocytes against oxidative stress-triggered damage and apoptosis, Eur. J. Pharmacol. (2007), doi:10.1016/j.ejphar.2007.04.031.

36. K. M. Brothers, R. L. Gratacap, S. E. Barker, Z. R. Newman, A. Norum, R. T. Wheeler, NADPH Oxidase-Driven Phagocyte Recruitment Controls Candida albicans Filamentous Growth and Prevents Mortality, PLoS Pathog. (2013), doi:10.1371/journal.ppat.1003634.

37. R. D. Gray, G. Hardisty, K. H. Regan, M. Smith, C. T. Robb, R. Duffin, A. MacKellar, J. M. Felton, L. Paemka, B. N. McCullagh, C. D. Lucas, D. A. Dorward, E. F. McKone, G. Cooke, Z. C. Donnelly, P. K. Singh, D. A. Stoltz, C. Haslett, P. B. McCray, M. K. B. Whyte, A. G. Rossi, D. J. Davidson, Delayed neutrophil apoptosis enhances NET formation in cystic fibrosis, Thorax (2018), doi:10.1136/thoraxjnl-2017-210134.

38. N. T. N. Trinh, O. Bardou, A. Privé, E. Maillé, D. Adam, S. Lingée, P. Ferraro, M. Y. Desrosiers, C. Coraux, E. Brochiero, Improvement of defective cystic fibrosis airway epithelial wound repair after CFTR rescue, Eur. Respir. J. (2012), doi:10.1183/09031936.00221711.

39. K. R. Schiller, P. J. Maniak, S. M. O’Grady, Cystic fibrosis transmembrane conductance regulator is involved in airway epithelial wound repair, Am. J. Physiol. Physiol. (2010), doi:10.1152/ajpcell.00215.2010.

40. P. Santabárbara-Ruiz, M. López-Santillán, I. Martínez-Rodríguez, A. Binagui-Casas, L. Pérez, M. Milán, M. Corominas, F. Serras, ROS-Induced JNK and p38 Signaling Is Required for Unpaired Cytokine Activation during Drosophila Regeneration, PLoS Genet. (2015), doi:10.1371/journal.pgen.1005595.

41. D. André-Lévigne, A. Modarressi, M. S. Pepper, B. Pittet-Cuénod, Reactive oxygen species and NOX enzymes are emerging as key players in cutaneous wound repair Int. J. Mol. Sci. (2017), doi:10.3390/ijms18102149.

42. D. J. McKeon, A. M. Condliffe, A. S. Cowburn, K. C. Cadwallader, N. Farahi, D. Bilton, E. R. Chilvers, Prolonged survival of neutrophils from patients with ΔF508 CFTR mutations Thorax (2008), doi:10.1136/thx.2008.096834.

43. R. Dinwiddie, Anti-inflammatory therapy in cystic fibrosis, J. Cyst. Fibros. (2005), doi:10.1016/j.jcf.2005.05.010.

44. C. Stumpf, Q. Fan, C. Hintermann, D. Raaz, I. Kurfürst, S. Losert, W. Pflederer, S. Achenbach, W. G. Daniel, C. D. Garlichs, Anti-inflammatory effects of Danshen on human vascular endothelial cells in culture, Am. J. Chin. Med. (2013), doi:10.1142/S0192415X13500729.

45. X. Chen, J. Guo, J. Bao, J. Lu, Y. Wang, The Anticancer Properties of Salvia Miltiorrhiza Bunge (Danshen): A Systematic Review, Med. Res. Rev. (2014), doi:10.1002/med.21304.

46. S. Gao, Z. Liu, H. Li, P. J. Little, P. Liu, S. Xu, Cardiovascular actions and therapeutic potential of tanshinone IIA Atherosclerosis (2012), doi:10.1016/j.atherosclerosis.2011.06.041.

47. C.-Y. Wu, J.-Y. Cherng, Y.-H. Yang, C.-L. Lin, F.-C. Kuan, Y.-Y. Lin, Y.-S. Lin, L.-H. Shu, Y.-C. Cheng, H. Te Liu, M.-C. Lu, J. Lung, P.-C. Chen, H. K. Lin, K.-D. Lee, Y.-H. Tsai, Danshen improves survival of patients with advanced lung cancer and targeting the relationship between macrophages and lung cancer cells, Oncotarget (2017), doi:10.18632/oncotarget.18767.

48. C. Hall, M. Flores, T. Storm, K. Crosier, P. Crosier, The zebrafish lysozyme C promoter drives myeloid-specific expression in transgenic fish, BMC Dev. Biol. (2007), doi:10.1186/1471-213X-7-42.

49. P. M. Elks, F. J. Van Eeden, G. Dixon, X. Wang, C. C. Reyes-Aldasoro, P. W. Ingham, M. K. B. Whyte, S. R. Walmsley, S. A. Renshaw, Activation of hypoxia-inducible factor-1α (hif-1α) delays inflammation resolution by reducing neutrophil apoptosis and reverse migration in a zebrafish inflammation model, Blood (2011), doi:10.1182/blood-2010-12-324186.

50. G. R. Holmes, S. R. Anderson, G. Dixon, A. L. Robertson, C. C. Reyes-Aldasoro, S. A. Billings, S. A. Renshaw, V. Kadirkamanathan, Repelled from the wound, or randomly dispersed? Reverse migration behaviour of neutrophils characterized by dynamic modelling, J. R. Soc. Interface (2012), doi:10.1098/rsif.2012.0542.

51. J. A. Lister, C. P. Robertson, T. Lepage, S. L. Johnson, D. W. Raible, nacre encodes a zebrafish microphthalmia-related protein that regulates neural-crest-derived pigment cell fate, Development (1999).

52. M. Sarris, J. B. Masson, D. Maurin, L. M. Van Der Aa, P. Boudinot, H. Lortat-Jacob, P. Herbomel, Inflammatory Chemokines Direct and Restrict Leukocyte Migration within Live Tissues as Glycan-Bound Gradients, Curr. Biol. (2012), doi:10.1016/j.cub.2012.11.018.

53. H. M. Isles, K. D. Herman, A. L. Robertson, C. A. Loynes, L. R. Prince, P. M. Elks, S. A. Renshaw, The CXCL12/CXCR4 Signaling Axis Retains Neutrophils at Inflammatory Sites in Zebrafish, Front. Immunol. (2019), doi:10.3389/fimmu.2019.01784.

54. A. Navis, L. Marjoram, M. Bagnat, Cftr controls lumen expansion and function of Kupffer’s vesicle in zebrafish, Development (2013), doi:10.1242/dev.091819.

55. V. Mugoni, A. Camporeale, M. M. Santoro, Analysis of oxidative stress in Zebrafish embryos, J. Vis. Exp. (2014), doi:10.3791/51328.

