## Supplementary material for "Deletion of *cftr* leads to an excessive neutrophilic response and defective tissue repair in a zebrafish model of sterile inflammation"

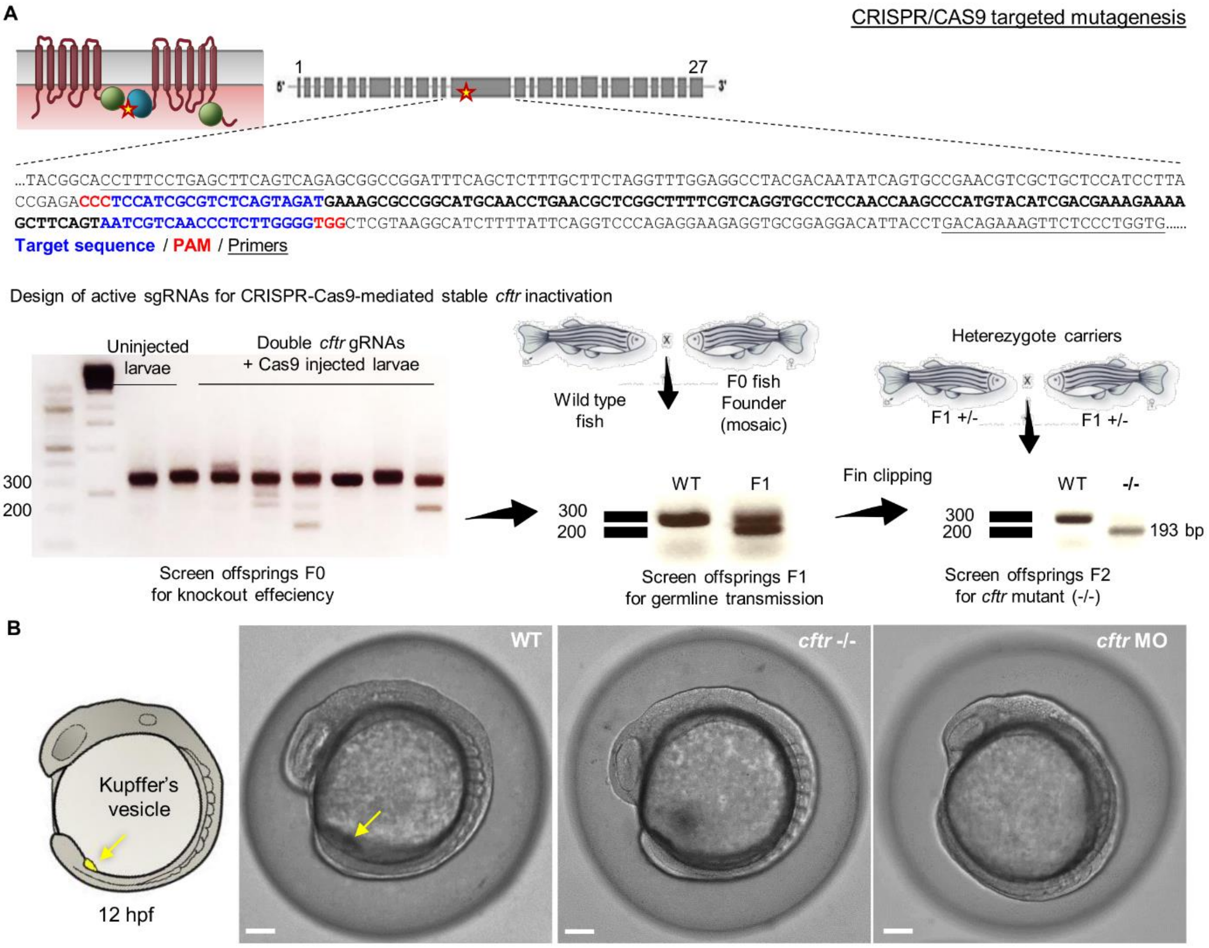


**Fig. S1. Generation of CFTR-depleted zebrafish**

(A) Generation of *cftr* null mutant in zebrafish using CRISPR-Cas9 gene editing. Schematic of the domain structure of CFTR and with a star indicating the CRISPR-Cas9 target site. Double guide RNA (gRNA) for CRISPR-Cas9-mediated stable *cftr* inactivation have been designed to effectively induce a deletion in *cftr* gene. The expected targeted deletion (127 bp) in F0 founder fish and germline transmission were detected and confirmed by PCR Analysis. Heterozygous carriers for the mutation are crossed to obtain *cftr* $^{-/-}$ homozygous mutant (#sh540). *cftr* -/- mutants are screening for impaired Kupffer's vesicle inflation at 8-somite stage.

(B) Loss of CFTR causes Kupffer's vesicle developmental defects in zebrafish. Representative photomicrography showing altered Kupffer's vesicle inflation in both *cftr* -/- mutant and *cftr* morphant.

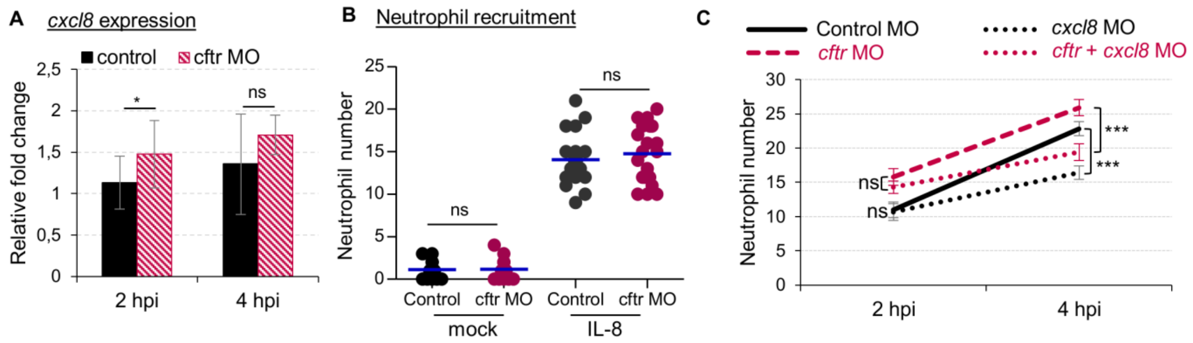


**Fig. S2. IL8-dependent neutrophil chemotaxis in *cftr*-defective zebrafish larvae**

(A) Tail transection was performed on control and *cftr* MO and mRNA levels of *cxcl8* gene determined by qRT-PCR in tail fin tissue at 2 and 4 hpi. Gene expression was normalized against *ef1a* and expressed as fold change over tail fin tissue from uninjured larvae (30 fins per replicat; mean relative ± SEM gene expression of 4 independent experiments; two-tailed Bonferroni t-test).

(B) Mean number of recruited neutrophils into the otic cavity in response to mock or IL8 injection in control and *cftr* MO *TgBAC(mpx:EGFP)i114 larvae* monitored at 2 hpi (n=20; means ± SEM from 3 independent experiments; two-tailed Bonferroni t-test). Cell counts in control or CFTR-depleted animal show no significant difference (ns) in the average number of neutrophils recruited at IL8-injected site.

(C) *TgBAC(mpx:EGFP)i114* controls, *cftr, Il8,* and double *cftr/il8* morphants were tail amputated and neutrophils at wounds were enumerated at 2 and 4 hpi (n=21; means ± SEM from 3 independent experiments; two-way ANOVA with Tukey post-test).

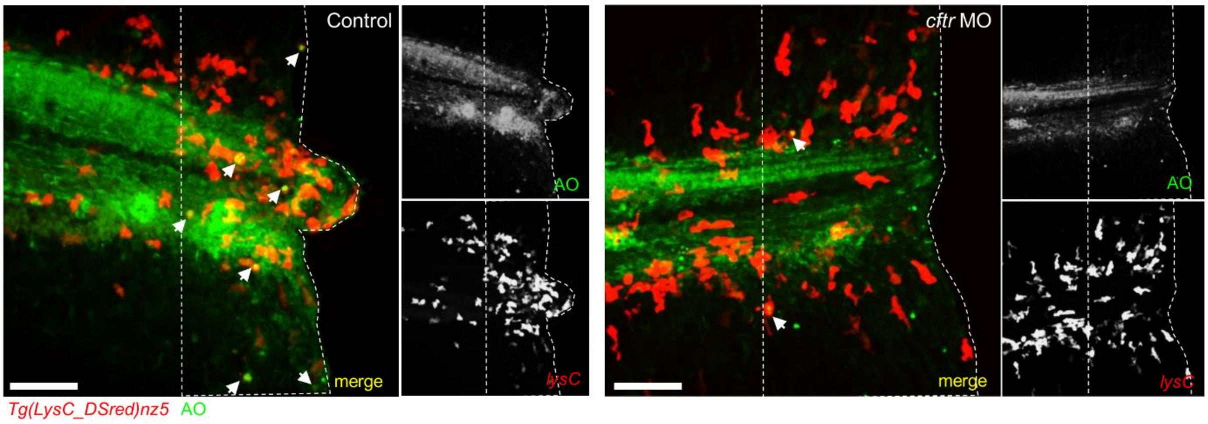


**Fig. S3. Neutrophil lifespan in *cftr*-defective zebrafish larvae**

3 dpf control and *cftr* MO *Tg(LysC:DSred)nz5* larvae were amputated and stained with acridine orange (OA) to label death cells. Representative confocal pictures of injured tails at 8 hpi (scale bars, 60 μm) revealing the proportion of dying neutrophils at the wound (white arrow). Dotted lines indicate the outline of the wounds.

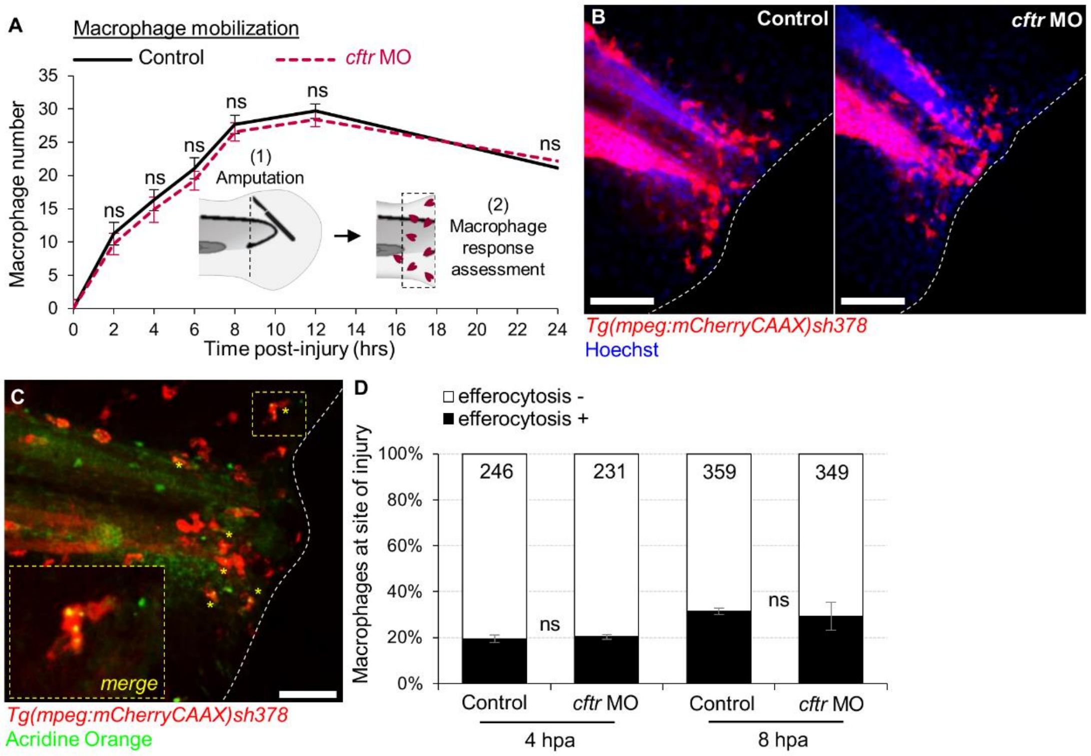


**Fig. S4. Macrophage behavior in *cftr*-defective zebrafish larvae**

(A-B) Macrophage recruitment assay in control and *cftr* MO *Tg(mpeg:mCherryCAAX)sh378* larvae 3 dpf larvae were tail injured and the number of macrophage mobilized at wound has been observed and recorded throughout inflammatory response by epifluorescence and confocal microscopies.

A) Dynamic of macrophage mobilization toward the wound. (n=30; mean ± SEM of 3 independent experiments; two-way ANOVA with Tukey post-test).

B) Illustrative confocal images of amputated tails in control and *cftr* morphant larvae showing the representative number of macrophages at site of injury at 12 hpi (scale bars, 200 μm).

(C-D) Macrophage efferocytosis assay in control and *cftr* MO *Tg(mpeg:mCherryCAAX)sh378* larvae. Animal were tail transected and stained with acridine orange (AO) for cell death visualization. Number of macrophages contained AO positive debris has been evaluated at 4 and 8 hpi at site of injury by confocal microscopy (scale bar, 100 μm).

C) Representative images of amputated tails at 8 hpi in control or *cftr* MO *Tg(mpeg:mCherryCAAX)sh378* stained with AO.

D) Macrophage efferocytosis rate at site of injury at 4 and 8 hpi. (n=21 larvae, mean ± SEM of 3 independent experiments, Fisher t test).

**Table 1. Primers used for qRT-PCR experiments**

| Genes | Sequences | References |
|---|---|---|
| *ef1a* | (F) 5′-TCTGTTACCTGGCAAAGGG-3’<br>(R) 5′-TTCAGTTTGTCCAACACCCA-3’ | *(1)* |
| *cxcl8* | (F) 5′-CCTGGCATTTCTGACCATCAT-3’<br>(R) 5′-GATCTCCTGTCCAGTTGTCAT-3’ | *(1)* |
| *duox2* | (F) 5′-GTTGGCTTTGGTGTAACTGTA-3’<br>(R) 5′-GCCCAGGCTGTGAGAG-3’ | *(2)* |

### Supplementary Materials and Methods

#### Morpholino injection

Morpholino were purchased from Gene Tools. The knockdown of *cftr* was carried on by injecting the validated *cftr* splice-blocking morpholino (5'-GACACATTTTGGACACTCACACCAA-3') into one-cell-stage zebrafish as previously described *(3)*. The morpholino for cxc*l8* knockdown (5'-TATTTATGCTTACTTGACAATGATC-3') *(4)* and *duox2* knockdown (5′-AGTGAATTAGAGAAATGCACCTTTT-3′) *(2)* were prepared and injected as described earlier. Neutrophil depleted-embryos were generated using the *csf3r* morpholino (5'-GAAGCACAAGCGAGACGGATGCCA*T-3'*) *(5)*. A standard control morpholino (5'-CCTCTTACCTCAGTTACAATTTATA-3') was used as a negative control.

#### Whole Body Neutrophil Counts

To assess total neutrophil number, 3 dpf *TgBAC(mpx*:*eGFP)i114* larvae were mounted in 0.8% agarose supplemented with tricaine (0.168 mg/mL; Sigma-Aldrich) and imaged on an Eclipse TE2000 U inverted compound fluorescence microscope (Nikon UK Ltd., Kingston upon Thames, UK) with a 4x NA objective lens. Three images were taken per larvae: one of the head region, one of the trunk region and one of the tail region. Neutrophils were counted manually, from these images and combined to give a whole body neutrophil count.

#### Neutrophils recruitment assays

The f-Met-Leu-Phe (fMLP, Sigma-Aldrich) or recombinant human IL-8 (rhIL-8, R&D Systems, Inc.) were used to induce neutrophil chemotaxis pharmacologically, as previously described *(3)*. Injured larvae were maintained at 28°C in sterile E3 media in a 24 well plate. Neutrophils at the injection sites were quantified and imaged on a fluorescence microscope (Nikon) using a 20x NA objective lens.

#### *In vivo* neutrophil reverse migration assay

Inflammation was elicited by tail transection on *Tg(mpx:gal4)sh267;Tg(UAS:kaede)i222* larvae following established protocols *(6)*. Briefly, injured larvae were raised to 4 hpi, then neutrophils at the wound site was photoconverted by 120 pulses of the 405-nm laser at 40% laser power using an UltraVIEW PhotoKinesis device on an UltraVIEW VoX spinning disk confocal microscope (PerkinElmer Life and Analytical Sciences). Larvae were transferred to an Eclipse TE2000-U inverted compound fluorescence microscope (Nikon), then were time-lapsed with 2.5min intervals between 5 and 9 hpi using a 1394 ORCA-ERA camera (Hamamatsu Photonics Inc).

Tracking analysis of red-fluorescing neutrophils moving away from photoconverted site was assessed in Volocity 6 (Improvision; PerkinElmer Life and Analytical Sciences), using the intensity of fluorescence to identify individually labeled neutrophils over time course of inflammation resolution.

**Neutrophil apoptosis assays**

Rates of apoptotic neutrophils were assessed in 4 and 8 hpi 4% paraformaldehyde-fixed larvae after dual staining with Rhodamine-TUNEL (ApopTag Red; Millipore Corp.) to label apoptotic cells and FITC-TSA (TSAplus kit; Fluorescence Systems, PerkinElmer Life and Analytical Sciences) to label neutrophils as previously described *(7)*.

Neutrophilic response at the wound were imaged by confocal microscopy (PerkinElmer Life), and the percentage of neutrophil apoptosis was calculated by comparing the total neutrophil number (TSA-positive) and number of apoptotic neutrophils (dual TSA/TUNEL-positive).

*In vivo cell death labeling and* **efferocytosis assays**

Rates of macrophage efferocytosis were assessed by staining with acridine orange (AO; Sigma-Aldrich) labeling death cells. 3 dpf *Tg(mpeg1:mCherry-CAAX)sh378* zebrafish larvae were tail-injured and AO-stained as described previously *(8)*.

Phagocytosis analysis of fluorescently labelled death cells was performed at 4 and 8 hpi by confocal microscopy (PerkinElmer Life), by comparing the number of macrophage phagocytosed AO-positive debris at the wound versus total mobilized macrophages.

*In vivo* **oxidative activity assay**

Oxidative response at the wound sites was measured using CellROX® Deep Red (ThermoFisher) following established protocols *(3)*. Larvae were incubated with CellROX® reagent prior tail fin amputation procedure, then immediately injured. ROS production at the wound was imaged by confocal microscopy (PerkinElmer Life) at 30 min. Oxidative response post injury at the wound was assessed in ImageJ 1.8 (National Institutes of Health) using the intensity of fluorescence and normalized to uninjured animal (WT or CF fish).

**qRT-PCR analysis**

Larvae were injured, then RNA from tails was extracted using TRIzol at time points indicated in Figure legends and cDNA synthesized using SuperScript II (Thermo Fisher Scientific). Real-time RT-PCRs were performed with a DyNAmo Flash SYBR green qPCR kit (Thermo Fisher Scientific) and gene expressions were detected with gene-specific primers listed in Table 1. Each experiment was run in triplicate. ΔCT was calculated using the housekeeping gene *ef1α* as a reference gene. Relative expression levels were calculated using the ΔΔCt method.

### References

1. A. Bernut, M. Nguyen-Chi, I. Halloum, J. L. Herrmann, G. Lutfalla, L. Kremer, Mycobacterium abscessus-Induced Granuloma Formation Is Strictly Dependent on TNF Signaling and Neutrophil Trafficking, *PLoS Pathog.* (2016), doi:10.1371/journal.ppat.1005986.

2. P. Niethammer, C. Grabher, A. T. Look, T. J. Mitchison, A tissue-scale gradient of hydrogen peroxide mediates rapid wound detection in zebrafish, *Nature* (2009), doi:10.1038/nature08119.

3. A. Bernut, C. Dupont, N. V. Ogryzko, A. Neyret, J. L. Herrmann, R. A. Floto, S. A. Renshaw, L. Kremer, CFTR Protects against Mycobacterium abscessus Infection by Fine-Tuning Host Oxidative Defenses, *Cell Rep.* (2019), doi:10.1016/j.celrep.2019.01.071.

4. M. Sarris, J. B. Masson, D. Maurin, L. M. Van Der Aa, P. Boudinot, H. Lortat-Jacob, P. Herbomel, Inflammatory Chemokines Direct and Restrict Leukocyte Migration within Live Tissues as Glycan-Bound Gradients, *Curr. Biol.* (2012), doi:10.1016/j.cub.2012.11.018.

5. N. Palha, F. Guivel-Benhassine, V. Briolat, G. Lutfalla, M. Sourisseau, F. Ellett, C. H. Wang, G. J. Lieschke, P. Herbomel, O. Schwartz, J. P. Levraud, Real-Time Whole-Body Visualization of Chikungunya Virus Infection and Host Interferon Response in Zebrafish, *PLoS Pathog.* (2013), doi:10.1371/journal.ppat.1003619.

6. C. A. Loynes, J. A. Lee, A. L. Robertson, M. J. G. Steel, F. Ellett, Y. Feng, B. D. Levy, M. K. B. Whyte, S. A. Renshaw, PGE2 production at sites of tissue injury promotes an anti-inflammatory neutrophil phenotype and determines the outcome of inflammation resolution in vivo, *Sci. Adv.* (2018), doi:10.1126/sciadv.aar8320.

7. P. M. Elks, F. J. Van Eeden, G. Dixon, X. Wang, C. C. Reyes-Aldasoro, P. W. Ingham, M. K. B. Whyte, S. R. Walmsley, S. A. Renshaw, Activation of hypoxia-inducible factor-1α (hif-1α) delays inflammation resolution by reducing neutrophil apoptosis and reverse migration in a zebrafish inflammation model, *Blood* (2011), doi:10.1182/blood-2010-12-324186.

8. A. Bernut, J.-L. Herrmann, K. Kissa, J.-F. Dubremetz, J.-L. Gaillard, G. Lutfalla, L. Kremer, Mycobacterium abscessus cording prevents phagocytosis and promotes abscess formation, *Proc. Natl. Acad. Sci.* (2014), doi:10.1073/pnas.1321390111.
